## Supplemental Figures for "Distinct beta frequencies reflect categorical decisions"

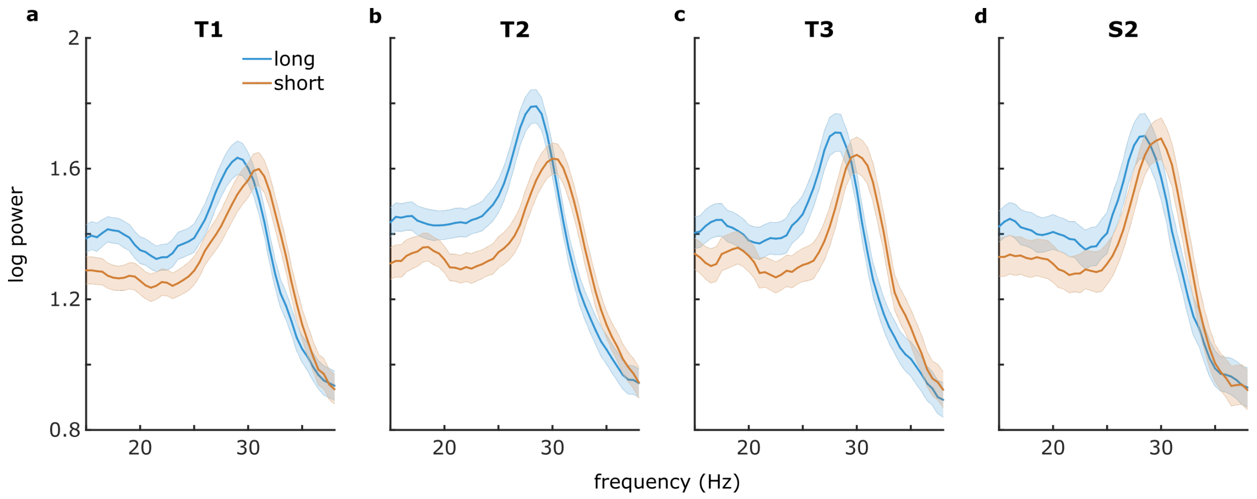


**Figure S1.** Beta peak frequency in monkey 1 dlPFC reflected the categorical decision during the decision delay in each version of the task. Power spectra for “long” stimulus (blue) vs. “short” stimulus trials (orange) during trials with correct responses **(a-c)** in the temporal categorization versions of the task with the shortest to longest stimuli respectively, and **(d)** the distance categorization task (see Figure 1b for exact values and Table 4 for statistics). Shaded regions around the line graphs represent the standard error of the mean.

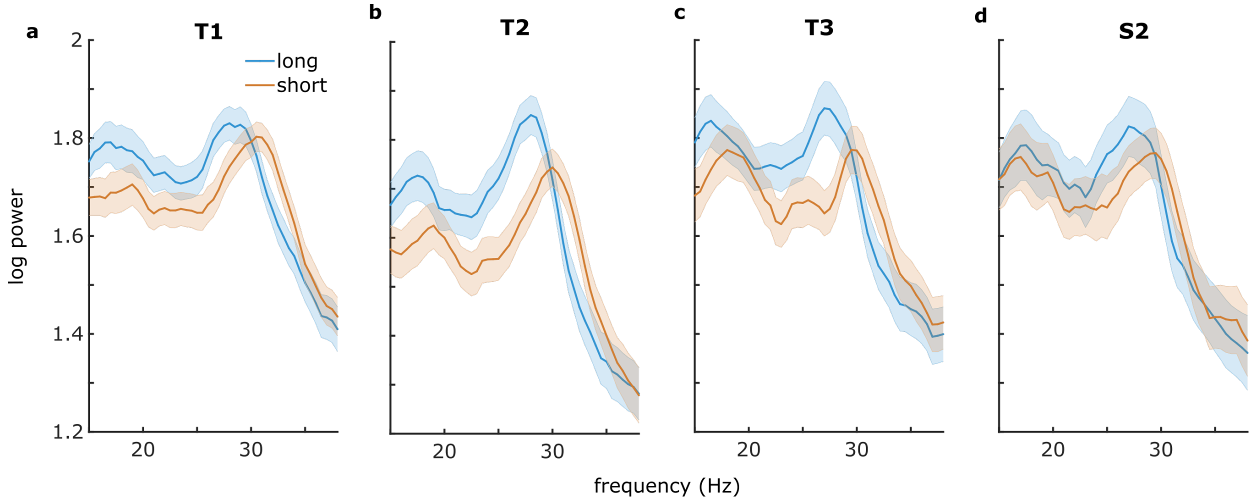


**Figure S2.** Beta peak frequency in monkey 1 preSMA reflected the categorical decision during the decision delay in each version of the task. Power spectra for “long” stimulus (blue) vs. “short” stimulus trials (orange) during trials with correct responses **(a-c)** in the temporal categorization versions of the task with the shortest to longest stimuli respectively, and **(d)** the distance categorization task (see Figure 1b for exact values and Table 5 for statistics). Shaded regions around the line graphs represent the standard error of the mean.

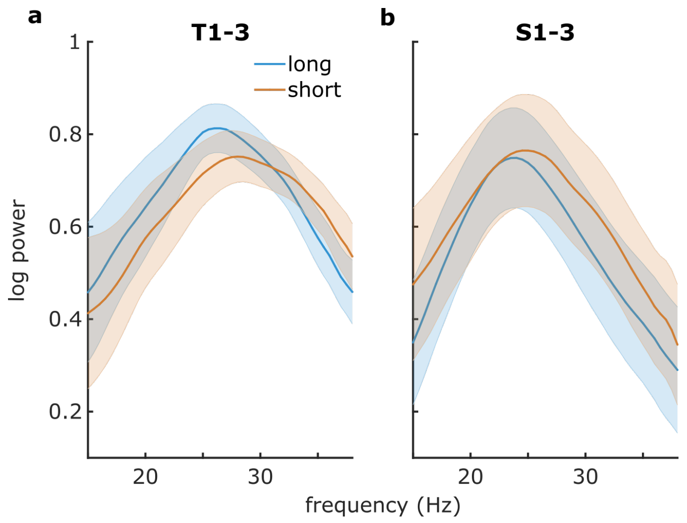


**Figure S3.** Beta peak frequency in monkey 2 preSMA reflected the categorical decision during the decision delay in each version of the task. Power spectra for “long” stimulus (blue) vs. “short” stimulus trials (orange) during trials with correct responses **(a)** in the temporal categorization versions of the task (pooled together) and **(d)** the distance categorization versions of the task (pooled together; see Figure 1b for exact stimulus values and Table 5 for statistics). Shaded regions around the line graphs represent the standard error of the mean.

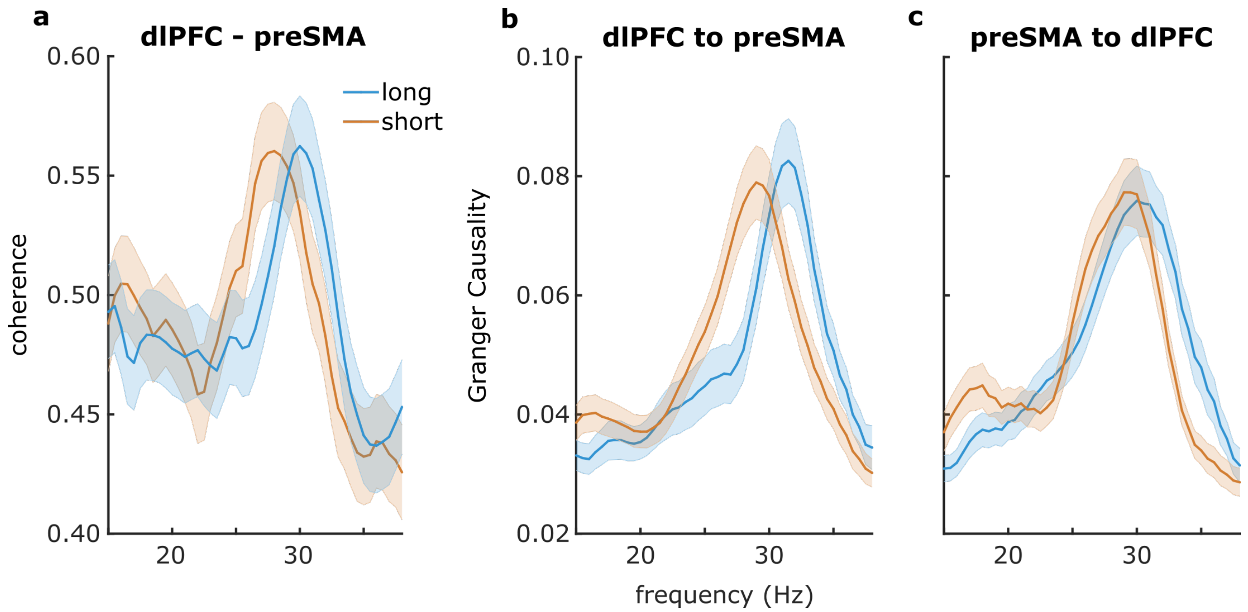


**Figure S4.** Peak frequencies of between-region connectivity reflected the categorical decision during the decision delay, even when the decision was incorrect. **(a)** Coherence between preSMA and dlPFC on incorrect trials. The direction of the peak frequency shift was reversed compared with correct trials (see Figure 4a). **(b, c)** Granger causality between preSMA and dlPFC. The direction of the peak frequency of dlPFC to preSMA and that of preSMA to dlPFC Granger causality were reversed compared with correct trials (see Figure 4 c, d). However, on both correct and incorrect trials, peak frequency of dlPFC to preSMA, but not preSMA to dlPFC Granger causality, reflected the categorical decision. Shaded regions around the line graphs represent the standard error of the mean.

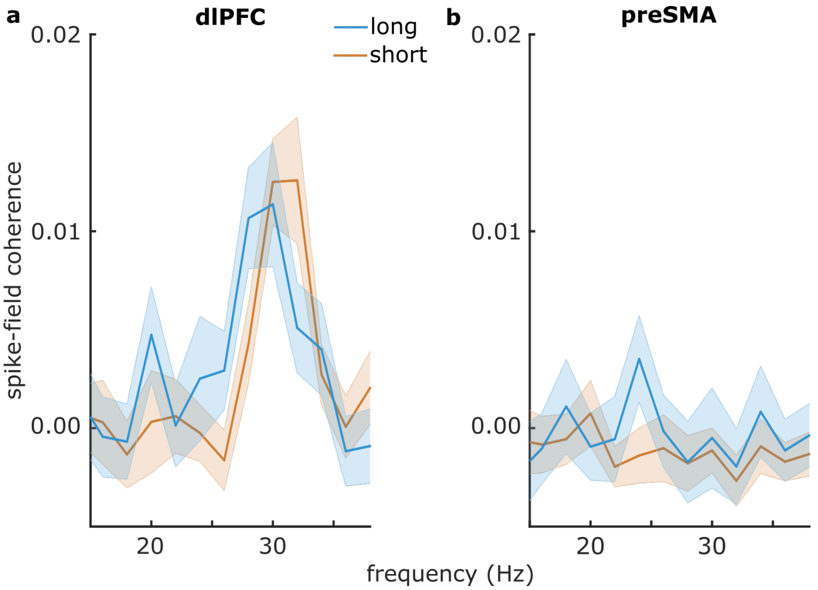


**Figure S5.** Spike-field coherence during the decision delay. **(a)** Spike-field coherence in dlPFC between short-selective neurons and the LFP during short-categorized trials, and between long-selective neurons and the LFP during long-categorized trials: the peak frequency reflected the categorical decision. **(b)** Same for preSMA: there were no significant peaks in the beta range.
