## Supplemental Tables for "Distinct beta frequencies reflect categorical decisions"

| **Monkey 1 dlPFC** | mean (+/- SD) prestimulus | mean (+/- SD) decision | decision > prestimulus (t) | decision > prestimulus (p) |
|---|---|---|---|---|
| frequency (Hz) | 24.8 (+/- 1.8) | 28.6 (+/- 2.3) | 23.3 | <1e-12 |
| burst frequency (Hz) | 22.3 (+/- 1.5) | 25.1 (+/- 2.3) | 14.4 | <1e-12 |
| burst number | 1.44 (+/- 0.23) | 1.76 (+/- 0.33) | 18.9 | <1e-12 |
| burst peak power (a.u.) | 3.71 (+/- 0.21) | 4.04 (+/- 0.31) | 15.9 | <1e-12 |
| burst time span (s) | 158 (+/- 19) | 194 (+/- 38) | 12.1 | <1e-12 |
| burst frequency span (Hz) | 6.35 (+/- 0.63) | 7.12 (+/- 0.96) | 10.1 | <1e-12 |

**Table 1.** Beta frequency shift and burst profile during prestimulus vs decision delay in Monkey 1 dlPFC. All t-value df = 198.

| **Monkey 1 preSMA** | mean (+/- SD) prestimulus | mean (+/- SD) decision | decision > prestimulus (t) | decision > prestimulus (p) |
|---|---|---|---|---|
| frequency (Hz) | 24.1 (+/- 1.5) | 26.8 (+/- 3.2) | 12.7 | <1e-12 |
| burst frequency (Hz) | 22.1 (+/- 1.3) | 23.7 (+/- 2.1) | 14.5 | <1e-12 |
| burst number | 1.62 (+/- 0.21) | 2.04 (+/- 0.31) | 30.3 | <1e-12 |
| burst peak power (a.u.) | 3.71 (+/- 0.24) | 4.13 (+/- 0.50) | 13.1 | <1e-12 |
| burst time span (s) | 170 (+/- 17) | 193 (+/- 27) | 10.8 | <1e-12 |
| burst frequency span (Hz) | 6.81 (+/- 0.58) | 7.31 (+/- 0.81) | 9.0 | <1e-12 |

**Table 2.** Beta frequency shift and burst profile of prestimulus vs. decision delay beta activity in Monkey 1 preSMA. All t-value df = 198.

| **Monkey 2 preSMA** | mean (+/- SD) prestimulus | mean (+/- SD) decision | decision > prestimulus (t) | decision > prestimulus (p) |
|---|---|---|---|---|
| frequency (Hz) | 21.0 (+/- 0.8) | 24.0 (+/- 1.9) | 6.8 | 3e-6 |
| burst frequency (Hz) | 21.4 (+/- 1.0) | 26.2 (+/- 2.2) | 12.8 | 4e-10 |
| burst number | 1.37 (+/- 0.22) | 1.06 (+/- 0.29) | -6.1 | 1e-5 |

| | | | | |
|---|---|---|---|---|
| burst peak power (a.u.) | 3.81 (+/- 0.20) | 4.43 (+/- 0.53) | 4.7 | 2e-4 |
| burst time span (ms) | 194 (+/- 19) | 127 (+/- 18) | -15.6 | 1e-11 |
| burst frequency span (Hz) | 6.63 (+/- 0.47) | 4.51 (+/- 1.05) | -7.3 | 1e-6 |

**Table 3.** Beta frequency shift and burst profile of prestimulus vs. decision delay beta activity in Monkey 2 preSMA. All t-value df = 17.

| **dlPFC** | short trials (Hz) mean (+/-SD) | long trials (Hz) mean (+/-SD) | short > long (T(df)) | short > long (p) |
|---|---|---|---|---|
| Monkey 1 - T1 | 30.0 (+/- 2.0) | 29.2 (+/- 1.4) | 2.6 (59) | .012 |
| Monkey 1 - T2 | 29.9 (+/- 1.8) | 28.0 (+/- 1.3) | 9.0 (56) | 2e-12 |
| Monkey 1 - T3 | 29.9 (+/- 1.7) | 28.0 (+/- 1.2) | 8.9 (44) | 2e-11 |
| Monkey 1 - S2 | 29.3 (+/- 1.4) | 28.2 (+/- 1.3) | 5.2 (36) | 8e-6 |

**Table 4.** Beta frequency shift in dlPFC during decision delay for trials categorized as “short” vs. “long”. T1, T2, and T3 are the interval categorization tasks with the shortest, middle, and longest sets of intervals (see Figure 1b for exact values), respectively. S2 is the distance categorization task with the middle set of distances (see Figure 1b for exact values).

| **preSMA** | short trials (Hz) mean (+/-SD) | long trials (Hz) mean (+/-SD) | short > long (t(df)) | short > long (p) |
|---|---|---|---|---|
| Monkey 1 - T1 | 30.1 (+/- 2.2) | 28.1 (+/- 2.0) | 5.4 (59) | 1e-6 |
| Monkey 1 - T2 | 29.3 (+/- 2.4) | 27.7 (+/- 1.8) | 4.2 (56) | 1e-4 |
| Monkey 1 - T3 | 29.0 (+/- 2.5) | 27.4 (+/- 1.6) | 3.3 (43) | .002 |
| Monkey 1 - S2 | 28.7 (+/- 2.0) | 27.3 (+/- 1.7) | 4.7 (35) | 4e-5 |
| Monkey 2 - all tasks | 25.2 (+/- 2.7) | 23.6 (+/- 2.8) | 2.7 (14) | .016 |

**Table 5.** Beta frequency shift in preSMA during decision delay for trials categorized as “short” vs. “long”. T1, T2, and T3 are the interval categorization tasks with the shortest, middle, and longest sets of intervals, respectively (see Figure 1b for exact values). S2 is the distance categorization task with the middle set of distances. For Monkey 2, tasks additionally included

S1 and S3, the distance categorization tasks with the shortest and longest distances, respectively.

| **Monkey 1 dlPFC** | mean (+/- SD) short trials | mean (+/- SD) long trials | short > long (t) | short > long (p) |
|---|---|---|---|---|
| burst frequency (Hz) | 28.7 (+/- 1.1) | 27.6 (+/- 1.0) | 10.8 | <1e-15 |
| burst number | 1.73 (+/-0.38) | 1.8 (+/- 0.34) | -2.1 | .039 |
| burst peak power (a.u.) | 3.95 (+/- 0.30) | 4.07 (+/- 0.39) | -3.4 | 6e-4 |
| burst time span (ms) | 194 (+/- 51) | 203 (+/- 45) | -1.8 | .070 |
| burst frequency span (Hz) | 6.84 (+/- 1.26) | 7.50 (+/- 1.10) | -5.5 | 8e-8 |

**Table 6.** Beta burst profile in dlPFC during decision delay for trials categorized as “short” vs. “long”. All t-value df = 198.

| **Monkey 1 preSMA** | mean (+/- SD) short trials | mean (+/- SD) long trials | short > long (t) | short > long (p) |
|---|---|---|---|---|
| burst frequency (Hz) | 28.4 (+/- 1.0) | 27.4 (+/- 0.9) | 9.9 | <1e-12 |
| burst number | 2.02 (+/- 0.33) | 2.05 (+/- 0.32) | -0.8 | .400 |
| burst peak power (a.u.) | 4.04 (+/- 0.43) | 4.19 (+/- 0.43) | -3.5 | 5e-4 |
| burst time span (ms) | 193 (+/- 37) | 202 (+/- 33) | -2.4 | 0.018 |
| burst frequency span (Hz) | 7.24 (+/- 1.07) | 7.69 (+/- 0.85) | -4.6 | 5e-6 |

**Table 7.** Beta burst profile in Monkey 1 preSMA during decision delay for trials categorized as “short” vs. “long”. All t-value df = 198.

| **Monkey 2 preSMA** | mean (+/- SD) short trials | mean (+/- SD) long trials | short > long (t) | short > long (p) |
|---|---|---|---|---|
| burst frequency (Hz) | 25.9 (+/- 1.5) | 25.2 (+/- 1.4) | 3.1 | .006 |
| burst number | 1.00 (+/- 0.29) | 0.97 (+/- 0.41) | -0.5 | .594 |
| burst peak power (a.u.) | 4.68 (+/- 0.71) | 4.59 (+/- 0.74) | 0.8 | .436 |

| | | | | |
|---|---|---|---|---|
| burst time span (ms) | 114 (+/- 15) | 128 (+/- 31) | -2.0 | .061 |
| burst frequency span (Hz) | 3.93 (+/- 1.23) | 4.37 (+/- 1.22) | -2.6 | .018 |

**Table 8.** Beta burst profile in Monkey 2 preSMA during decision delay for trials categorized as "short" vs. "long". All t-value df = 17.

| **Monkey 1 dlPFC** | long trials (Hz) mean (+/- SD) | short trials (Hz) mean (+/- SD) | short > long (t (df)) | short > long (p) |
|---|---|---|---|---|
| 450ms | 29.1 (+/- 1.9) | 30.2 (+/- 2.1) | 2.6 (99) | .011 |
| 500ms | 29.1 (+/- 1.6) | 30.1 (+/- 2.0) | 2.9 (99) | .005 |
| 870ms | 28.5 (+/- 1.6) | 30.1 (+/- 1.7) | 4.4 (81) | 4e-5 |
| 920ms | 28.9 (+/- 2.0) | 30.5 (+/- 1.5) | 6.42 (82) | 8e-9 |

**Table 9.** Beta frequency shift in dlPFC during decision delay of the duration categorization tasks, for trials with identical stimuli within different task versions (i.e., trials with identical stimuli categorized as "short" in one task version but "long" in another; see overlapping stimuli outlined in Figure 1b).

| **Monkey 1 preSMA** | long trials (Hz) mean (+/- SD) | short trials (Hz) mean (+/- SD) | short > long (t (df)) | short > long (p) |
|---|---|---|---|---|
| 450ms | 29.2 (+/- 2.5) | 30.1 (+/- 2.1) | 1.99 (99) | .049 |
| 500ms | 28.9 (+/- 2.0) | 30.0 (+/- 2.3) | 2.8 (99) | .006 |
| 870ms | 28.2 (+/- 1.6) | 29.3 (+/-2.3) | 2.8 (81) | .007 |
| 920ms | 28.4 (+/- 1.7) | 29.6 (+/- 2.0) | 2.9 (82) | .005 |

**Table 10.** Beta frequency shift in preSMA during decision delay of the duration categorization tasks, for trials with identical stimuli within different task versions (i.e., trials with identical stimuli categorized as "short" in one task version but "long" in another; see overlapping stimuli outlibed in Figure 1b).
